# Female acoustic signaling of sexual immaturity depresses male courtship in Drosophila

**DOI:** 10.64898/2026.06.09.731064

**Authors:** Martin Bernet, Anne C. von Philipsborn

## Abstract

In an early phase of life, most animals are behaviorally and physiologically not yet able to reproduce but show adaptations specific to a juvenile state. In Drosophila females, sexual maturation after metamorphosis is achieved by acquiring receptivity to male courtship and completing oogenesis, a transition that is under hormonal control and requires coordinates changes in the nervous system. Here, we show that immature females display a transient signaling behavior during the time window of asexuality by flicking their wings in response to and dependent on male courtship stimuli. Immature wing flicks require the activity of Doublesex (Dsx) expressing central brain pC1a neurons that mediate receptivity in mature virgins. Immature wing flicks generate patterned sound pulse trains that differ from other intraspecific acoustic signals, but resemble pulses produced during mature male agonistic interactions. Courting males exposed to immature flicks shorten courtship singing and abstain from copulation attempts, indicating that immature wing flicks serve as an effective rejection signal of asexual females to minimize futile male mating pursuits.

## Introduction

For successful sexual reproduction, animals need to choose a mate, but also an optimal time point during their life history. Multicellular animals generally undergo a period of sexual maturation characterized by growth, differentiation and changes in their behavioral repertoire before they can reproduce with full fecundity. In both mammals and insects, sexual maturation occurs under the influence of hormones that affect reproductive organs and the nervous system (Arnold 2009, Herbison 2016, Santos et al. 2019, Truman and Riddiford 2023). In mammals, the core circuit elements for reproductive behavior are established in the nervous system already in an early organizational phase before or around birth, to further reorganize and become fully functional in puberty (McCarthy and Arnold 2011, Schulz and Sisk 2016).

Sex differences in the Drosophila nervous system arise during pupal stages largely through selective neuronal apoptosis and differential axonal pathfinding (Kimura 2011, Sato and Yamamoto 2020, Goodwin and Hobert 2021, Allen et al. 2025). In the adult stage, these sexual dimorphisms in neuronal number, morphology and connectivity specify behavioral differences between the sexes (Asahina 2018, Meeh et al. 2021, Tong et al. 2026). Males respond to females with courtship and show male-specific patterns of aggression when competing among each other for resources. Virgin females accept males for copulation when courted appropriately, while mated females are less receptive and lay eggs on suitable oviposition sites. While largely innate, sexual behavior also shows some flexibility and is subjected to modulation depending on internal state, external conditions and experience (Ellendersen and von Philipsborn 2017, Karingo and Deutsch 2022, Day and Rezaval 2026). Besides pheromones sensed by olfaction or gustation, acoustic signals generated by precisely timed wing movements play an important role in Drosophila intraspecific communication during mating and aggression. Male courtship song, a close-range signal with a species-specific pattern, stimulates female receptivity. Females produce a distinct song during copula, that reflects the composition of transferred inseminate. During male-male aggressive encounters, a third type of wing sounds modulates the agonistic interactions. Neuronal circuits in the male and female nervous system are specialized to produce, detect and evaluate these signals and respond to them with specific behaviors (Baker et al. 2019, Swain and von Philipsborn 2021).

When flies emerge from the pupal stage at eclosion, they are not yet competent to perform all adult behaviors. In a period of maturation, they undergo a series of coordinated physiological and behavioral changes. The most striking among these is the transition to sexuality, which corresponds to the appearance of courtship behavior in the male and the development of receptivity and the ability to accept copulations in females (Ford et al. 1989, White and Ewers 2014, Seong et al. 2023, Yip et al. 2024).

In male flies, the mechanisms underlying the transition from an asexual to a reproductive state are still under debate (Zhang et al. 2021, Ji et al. 2023), with evidence for organizational changes in the connectivity and excitability of courtship brain circuits (Ji et al. 2023). Juvenile Hormone (JH) promotes male mating behavior (Wilson et al. 2003, Wijesekera et al. 2016, Lin et al. 2016), growth of the reproductive tract (Box et al. 2024, Ramesh et al. 2025), the production of seminal fluid (Herndon et al. 1997) and fecundity (Meiselman et al. 2017).

In immature females, the rapid “switch-on” of receptivity during the first day after eclosion depends on the action of JH (Ringo et al. 1991, Ringo et al. 2005, Bilen et al. 2013). JH titers decrease in aging virgins and increase again after mating (Bownes and Rembold 1987, Moshitzky et al. 1996, Sugime et al. 2017). Elevated JH titers after mating stimulate oogenesis (Bownes 1982, Soller et al. 1999, Zhang et al. 2022), but they are not, as during maturation, accompanied by an increase in receptivity. In contrast, mating leads to a decrease of receptivity, that is triggered by mechanical stimulation during copulation and the receipt of male inseminate containing sex peptide (Liu and Kubli 2003, Shao et al. 2019). Receptivity of mated females remains low for up to two weeks, as long as sperm and sex peptide are present in sperm storage organs (Peng et al. 2005, Singh et al. 2018) and sex peptide signals are relayed to the receptivity circuits in the female central nervous system (Yapici et al. 2008, Häsemeyer et al. 2009, Yang et al. 2009, Feng et al. 2014, Wang et al. 2021).

The lack of receptivity in immature versus mated females is thus mechanistically distinct. Early studies of Drosophila sexual behavior already revealed a corresponding behavioral difference in how immature and mated females respond to male courtship: while immature females flick their wings, mature females extrude their ovipositor (Manning 1967, Ewing and Bennet-Clark 1968, Connolly and Cook 1971). Ovipositor extrusion and its neural control have been recently studied in detail, suggesting that mating-induced ovulation gates the expression of this rejection response (Mezzera et al. 2020, Wang et al. 2020a). Much less is known about immature wing flicking, a behavior that stands out as being specific for the immature state that is otherwise mostly characterized by the lack of behavioral elements and attenuated response to stimuli. It has been anecdotally observed that immature wing flicks produce acoustic pulses (Pailette et al. 1991, Swain and von Philipsborn 2021). Otherwise, immature wing flicking has so far not been characterized regarding its development, proximate causation, neuronal mechanism and behavioral function.

Here, we fill this gap and characterize immature wing flicking behavior and its relationship to sexual maturation and the development of female receptivity.

We find that immature wing flicking produces structured acoustic pulse trains that are distinct from other wing signals occurring during intraspecific communication and resemble most closely the pulse trains produced by male-male agonistic interactions. Wing flicking occurs exclusively in a short time window after eclosion and is negatively regulated by JH signaling. Immature females flick their wings predominately shortly after male courtship song bouts. Both acoustic and pheromonal courtship stimuli from the male are required to fully trigger immature wing flicking. Immature wing flicking depends on central pC1a neurons that control receptivity in mature females. Wing flicking is a reliable signal of asexuality, highly efficient in depressing male copulation attempts. In this capacity, immature wing flicks differ from the ovipositor extrusion, the characteristic rejection response of mature mated females and function similarly to agonistic wing flicks of mature males.

## Results

### Immature females produce wing flick sounds when courted by males

To assess the acoustic behavior of immature virgins, we collected female flies immediately after eclosion and exposed them to mature males at 5 h later. In audio recordings of such pairings, we detect pulses that are clearly distinct from male courtship song and are absent when males court mature females. The immature female pulses resemble the ones produced by mature males during agonistic encounters (**Figure 1A**). High speed video recordings with a framerate of 500 Hz reveal that immature females perform rapid wing flicks corresponding to these pulses. Females extend both wings for the duration of a wing flick, with an angle of 77 ±3° (mean± sem, n=13 flicks from 6 flies) (**Figure 1B**). Each immature female wing flick pulse consists of multiple cycles and has a duration of 20-25 ms (**Figure 1A**). Wing flick pulses appear in trains, with 70% of inter pulse intervals (IPI) shorter than 300 ms (**Figure 1C**). The comparison of key acoustic properties such as pulse train length and of immature female wing flicks with wing sounds produced by males shows that immature female wing flicks resemble agonistic wing flicks of mature males more closely than male courtship pulse song but have a lower median pulse carrier frequency (**Figure 1D**). Immature wing flick sounds are also distinct from the copulation song of mature females, which consists of regular polycyclic pulses of higher frequency (Kerwin et al. 2020, Swain and von Philipsborn 2021).

**Figure 1.**
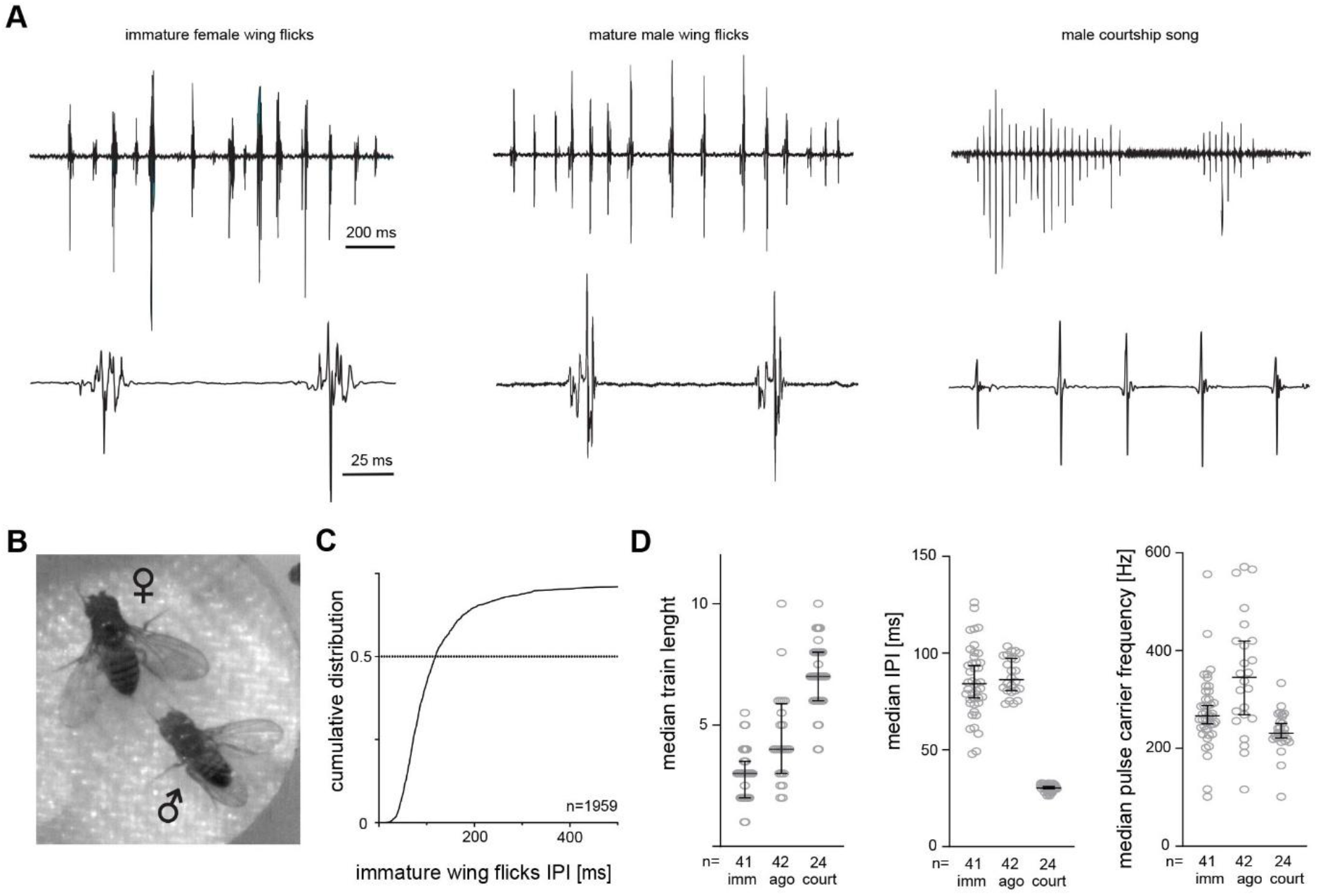
Immature females produce wing flick sounds when courted. **A** Oscillograms of immature female wing flick sounds, mature male wing flicks during agonistic encounters and male courtship song. Scale bars: 200 ms and 25 ms. **B** Video still from a high-speed recording of a mature male paired with a 5 h old virgin female, illustrating unilateral male singing posture and bilateral wing extension during immature female wing flicking. **C** Cumulative distribution of IPI length from immature wing flick sound pulses, n= 1959 IPIs from 10 min recordings of n=41 immature females paired with mature males. **D** Median pulse train length, IPI and pulse carrier frequency of immature female wing flicks (imm), agonistic wing flicks between mature males (ago) and courtship song of mature males (court). Each datapoint represents data from a 10 min recording of fly pairs (imm, court) or triads (ago), error bars indicate median and interquartile range.

### Immature wing flicks depend on female maturation state

To understand how female wing flicks are correlated with the maturation process, we paired female virgins at different time points after eclosion with mature males and recorded their sound production. Female immature wing flicks are seen at 2, 5 and 12 h after eclosion and drop to a very low level at 24 h that is undistinguishable from the level seen in mature mated females. The occurrence of immature female wing flicks is most prominent at 5h after eclosion, with virgins of this age producing significantly more flicks than at 2, 12 or 24h after eclosion (**Figure 2A**). We next wondered how immature flicks related to receptivity to mating. To observe mating during the first 1-2 days after eclosion in an undisturbed manner, we placed a female pupa with a mature male in a food vial and videotaped the interactions of the pair after female emergence. Mating was never observed before 10h after eclosion (**Figure 2B**). We conclude that immature females flick their wings most pronouncedly when they are not receptive to mating. Shortly after eclosion, flies deploy their compactly folded wings (White and Ewers 2014, Hadjaje et al. 2024) and the cuticle undergoes hardening to support effective wing movements. To compare wing flick production with flight ability during the first day after eclosion, we performed forced flight tests. At 2 h after eclosion, over 50% of flies are unable to fly. Flight performance improves significantly at 5 h, although it is still not at the optimum at 24 h (**Figure 2C**). Most wing flicking is thus produced when an intermediate flight ability has developed, but females do not yet engage in copulations.

**Figure 2.**
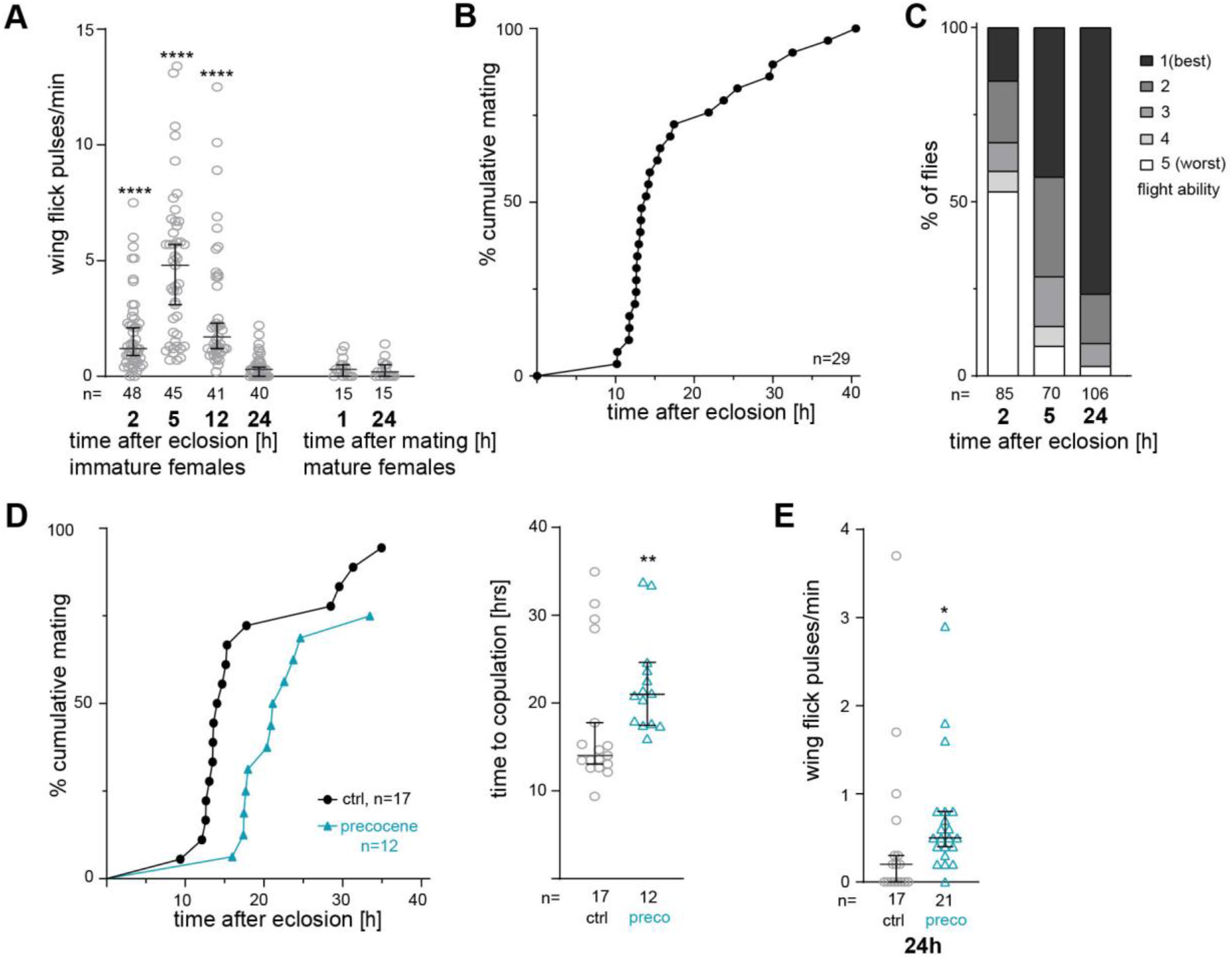
Immature wing flicks are specific for a short time window during maturation. **A** Amount of wing flicks in pulses/min produced by immature and mature females courted by males at different times after eclosion and mating, respectively. **** p<0.001, Kruskal-Wallis test with Dunn’s multiple comparison test, comparing flicks at 2, 5 and 12 hrs after eclosion to flicks at 24 hrs after eclosion. **B** Cumulative mating rate of females after eclosion, tested individually paired with a mature male in a food vial under constant observation. **C** Flight ability assessed by forced flight test of female flies at 2, 5 and 24 hrs after eclosion. **D** Cumulative mating rate of females after eclosion (left) and time to copulation after eclosion (right), with exposure to the antijuvenoid precocene and acteon control, respectively. **p=0.009, Mann-Whitney test. **E** Amount of wing flicks in pulses/min produced at 24 hrs after eclosion by females exposed to precocene and acteon control, respectively. * p=0.0103, Mann-Whitney test. In A, E, each datapoint represents data from a 10 min recording of one female paired with a courting mature male, error bars indicate median and interquartile range.

Reproductive maturation and the timing of the first copulation in Drosophila females is under the control of juvenile hormone (JH) signaling (Ringo et al. 2005, Bilen et al. 2013). To test if immature wing flicks were influenced by JH signaling as well, we treated females after eclosion with the antijuvenoid precocene I. As expected, blocking of JH synthesis delays the onset of mating after eclosion (**Figure 2D**). Concomitantly, females that were delayed in maturation by precocene I treatment show still significant amounts of wing flick behavior at 24h after eclosion, a time point when most control females have stopped to do so (**Figure 2E**).

We conclude that immature females courted by mature males show wing flicking that is accompanied by characteristic sound pulses. Wing flicks only occur within the first 12 h after eclosion, with a peak around 5 h, when most individuals have acquired the ability to use their wings for flight, but are not receptive for mating. Delaying reproductive maturation by interference with JH signaling prolongs the time period during which immature females produce wing flicks.

### Immature females wing flicks depend on multiple aspects of male courtship

We next asked which sensory stimuli could trigger immature wing flicks and to which extent wing flicking depended on courtship cues. For this, we paired immature females with female and male flies of different age. While pairing an immature female with a mature male elicits marked flicking, the presence of a mature female or an immature individual of either sex does not provoke wing flicks above the low background level observed in solitary immature females (**Figure 3A**). Mature males vigorously courted the immature females in our experiments. In the oscillograms of audio recordings, immature wing flick trains were often observed shortly after the onset of a male pulse song train (**Figure 3B**). To characterize the temporal relationship between the wing sounds of the two sexes in more detail, we measured the latency between the start of each male pulse song train and the closest immature wing flick following afterward for 42 pairs of flies. Immature wing flicks occur with significantly higher probability within 100-600 ms after the onset of male pulse song trains, peaking at the latency window of 300-400 ms. For this latency window, immature females thus listen on average to 10 male pulses before producing a wing flick (**Figure 3C**). Given the correlation between male courtship song and immature wing flicks, we wondered if immature flicks depended on male song or on other components of male courtship, such as general courtship activity (following, tapping) and pheromone profile. To disentangle which properties influenced immature flicking, we tested the effect of courtship song, courtship steps other than singing and pheromones by different experimental manipulations of males or females. Males lacking the male specific isoform of *fruitless* (*fruF* mutants) were used as non-courting animals with male specific pheromone profile, and females expressing the male specific isoform of *fruitless* (*fruM* mutants) were used as animals performing male courtship but lack the male pheromone profile (Demir and Dickson 2005). Removal of acoustic stimuli from male courtship song by muting courting males via wing amputation or deafening immature females via arista amputation reduces immature flicks, as does the complete removal of all courtship steps in presence of *fruF* mutant male. Removal of male specific pheromonal stimuli, either by using anosmic immature females (*IR8a, IR25a, GR63a, Orco* quadruple mutants, Ramdya et al. 2014) as focal animals or by using *fruM* mutant females as courting partners, likewise reduces immature flicks. Absence of one male stimulus (courtship song/behavior or pheromones), however, was not sufficient to completely abolish immature wing flicks to a level observed in the presence of mature males (**Figure 3D**). These findings indicate that different components of male courtship contribute to eliciting immature wing flicks, with male song, courtship following and male pheromones necessary for the full occurrence of the behavior.

**Figure 3.**
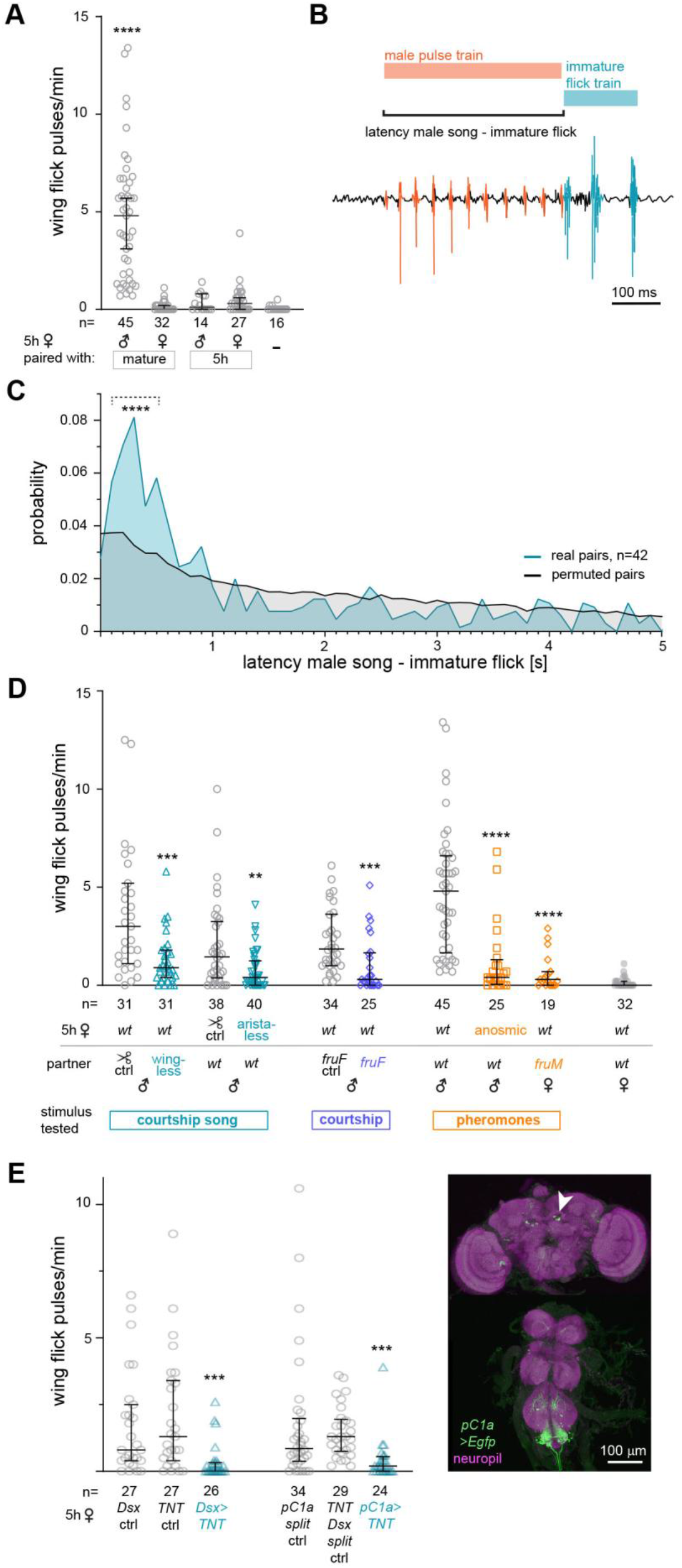
Immature wing flicks depend on male courtship signals and the female neurons processing them. **A** Amount of wing flicks in pulses/min produced by immature females exposed to partner flies with differing courtship activity (mature and immature males and females). **** p<0.0001, Kruskal-Wallis test with Dunn’s multiple comparison test, comparing flicks of paired flies with flicks of solitary females. **B** Example oscillogram showing a common temporal relationship of a male courtship song pulse train (orange) and an immature female wing flick train (teal), with a latency of the start of the male song train and the immature flick train of 350 ms. Scale bar: 100 ms. **C** Probability of immature wing flick trains occurring after a given latency (binned in 100 ms intervals) of the start of a male courtship train. Data from 42 pairs of immature females with mature males (recorded for 10 min each, with 653 male song-female flick events evaluated) is shown in teal, mean data from 1000 permuted datasets with misassigned male-female pairs is shown in black. Dashed bracket indicates the 5 100 ms latency bins (100 ms-600 ms) with significant enrichment of immature flick occurrence (**** p=0 for peak latency bin 300-400 ms, permutation testing). **D** Amount of wing flicks in pulses/min produced by immature females while different sensory stimuli of male courtship are experimentally manipulated. In teal removal of acoustic stimuli by male wing or female arista amputation, in blue removal of courtship by male *fruF* mutation, in orange removal of pheromone stimuli by quadruple smellblind mutations in females (anosmic) or exposure to a *fruM* mutant female courtship partner. ** p=0.0036, *** p=0.003, **** p<0.0001, Mann-Whitney tests. **E** Amount of wing flicks in pulses/min produced by immature females with Dsx+ neurons (*Dsx>TNT*) or pC1a (*pC1a>TNT*) neurons silenced by Tetanus toxin and respective genetic controls. *** p<0.001, Kruskal-Wallis test with Dunn’s multiple comparison test. Expression pattern of pC1a split GAL4 driver line (green) in the central nervous system of a 5 h old female, neuropil staining (anti-Bruchpilot) in magenta. Arrowhead indicates the three somata of the pC1a neuronal class on one side of the posterior brain. Scale bar: 100 μm. In A, D, E, each datapoint represents data from a 10 min recording of one female paired with the indicated partner, error bars indicate median and interquartile range.

### Female Doublesex expressing neurons are required for immature flicks

Given the finding that immature female flicking behavior depends on courtship stimuli from mature males, we wondered if it required the neuronal circuits implicated in processing male stimuli in mature females. The response of mature virgins to male courtship depends on neurons expressing the sex determination factor Doublesex (Dsx) (Zhou et al. 2014). Neuronal silencing of all Dsx positive (Dsx+) neurons by GAL4 UAS mediated expression of Tetanus toxin (TNT), which blocks synaptic transmission, leads to a loss of wing flicking behavior in immature females (**Figure 3E**). Among Dsx+ neurons, the class of pC1 neurons in the brain has been proposed as a prominent integration center of courtship stimuli (Zhou et al. 2014). pC1 contains different subtypes, pC1a-e, that are functionally distinct, with pC1a-c controlling receptivity and pC1d-e inter-female aggression (Schretter et al. 2020, Wang et al. 2020b, Wang et al. 2021). Using a GAL4 driver specific for pC1a (Yun et al. 2024), we find that silencing the synaptic activity of this subtype abolishes immature wing flicks. In immature females, this driver line (*R52G04-AD, dsx-DBD*) labels three cells of the pC1 cluster on each side of the brain (**Figure 3E**). This suggests that the same Dsx+ pC1a neuronal subtype that is required for mature virgin receptivity is also required for wing flicking behavior in immature virgins.

### Immature wing flicks shorten male song trains and suppress futile copulation attempts

Immature flicks are most prominent when females do not accept copulation and are stimulated by male courtship, suggesting the possibility that they act as rejection behavior and signal to the male that the female will not accept copulation. To probe this hypothesis, we asked how female flicks affected male courtship behavior. We first assessed male courtship song toward two types of immature females unable to perform wing flicks: wing amputated females or females with silenced Dsx+ neurons. In both cases, the total amount of pulse song the male sings toward the female is not changed (**Figure 4A**). However, the mean length of male pulse song trains is longer when the male courts immature females that do not produce flicks (**Figure 4B**). This indicates that immature wing flicks occurring shortly after the onset of male trains interrupt and stop male pulse song. Next, we tested the occurrence of male copulations attempts. Immature females unable to produce wing flicks in response to male courtship are exposed to an increased number of copulation attempts (**Figure 4C**). To test if wingless females were inherently more attractive to males and elicited more copulation attempts, we also tested wing amputated versus intact mated females. Mature mated females, which never produce wing flicks when courted, elicit the same number of copulation attempts irrespective of the presence of wings (**Figure 4D**). We conclude that immature female wing flicks suppress male copulation attempts. Of note, the increased number of copulation attempts toward immature virgins in the absence of wing flicks does not result in successful copulations. None of the immature 5 h old females without wing flick behavior (wing amputated or Dsx+ neuron silenced) that we tested in assays probing courtship song or copulation attempts (total n=112) accepted the partner male for copulation within the 10 min test time. Consistently, there is no advance in the onset of mating of wingless virgins compared to intact control flies, when they are tested continuously paired with a mature male on food during the first day after eclosion (**Figure 4E**). We therefore conclude that wing flicks do not function in preventing copulation but rather act to inhibit copulation attempts that are predetermined to be futile.

**Figure 4.**
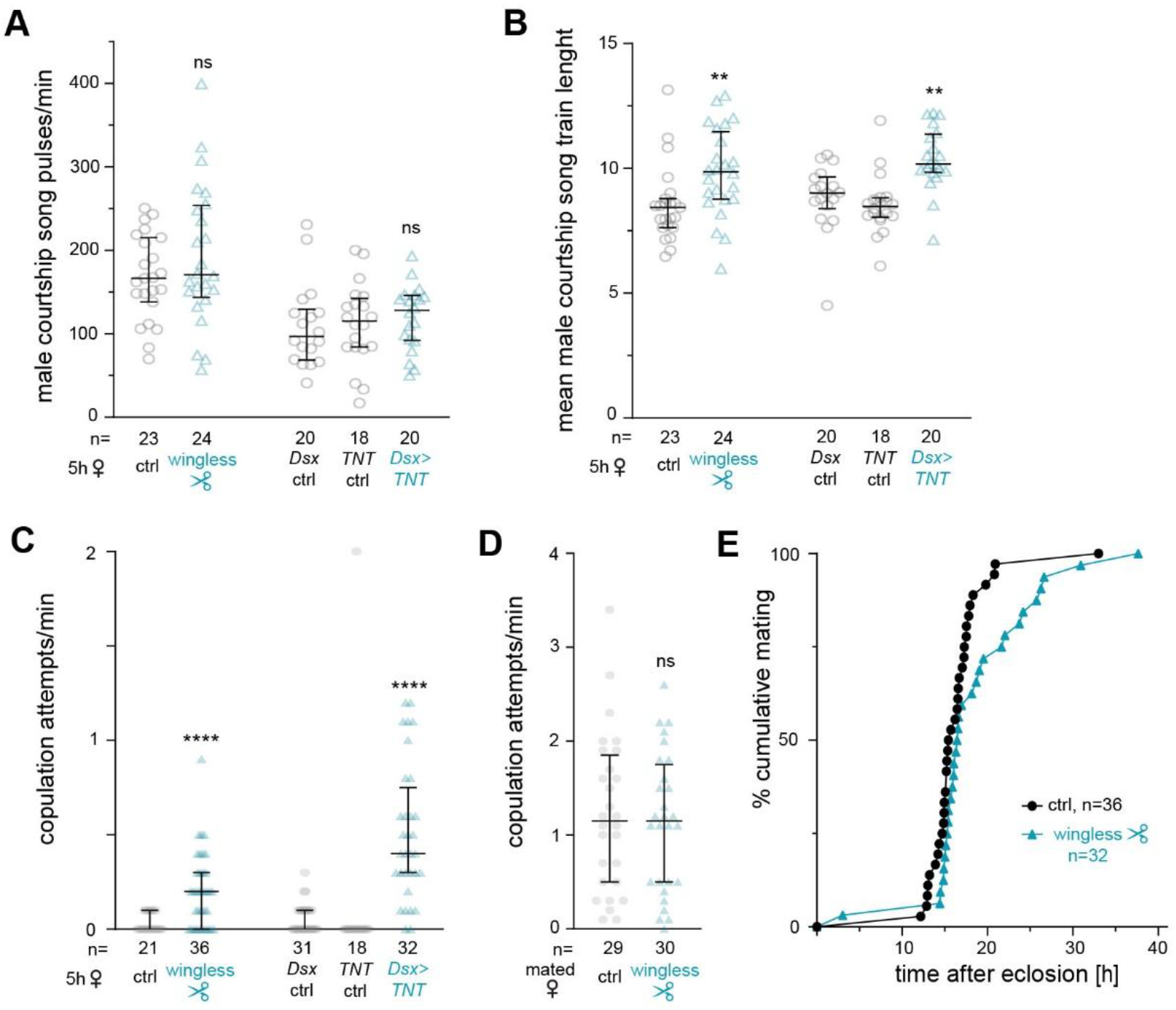
Immature wing flicks depress male courtship intensity. **A** Total amount of courtship pulse song produced by mature males courting immature females unable to produce wing flicks (wingless by amputation and with Dsx+ neurons silenced by Tetanus toxin, *Dsx>TNT*) and respective controls. ns, not significant, Mann Whitney test (ctrl vs wingless) Kruskal-Wallis with Dunn’s multiple comparison test (*Dsx>TNT* vs ctrls) **B** Mean male courtship pulse song train length of mature males courting immature females unable to produce wing flicks (wingless by amputation and with Dsx+ neurons silenced by Tetanus toxin, *Dsx>TNT*) and respective controls. **p=0.0019, Mann Whitney test (ctrl vs wingless) and **p=0.0057 Kruskal-Wallis with Dunn’s multiple comparison test (*Dsx>TNT* vs ctrls). **C** Frequency of copulation attempts of mature males courting immature females unable to produce wing flicks (wingless by amputation and with Dsx+ neurons silenced by Tetanus toxin, *Dsx>TNT*) and respective controls. ****p<0.0001, Mann Whitney test (ctrl vs wingless) and Kruskal-Wallis with Dunn’s multiple comparison test (*Dsx>TNT* vs ctrls). **D** Frequency of copulation attempts of mature males courting mature mated females, intact and wingless by amputation. ns, not significant, Mann Whitney test. **E** Cumulative mating rate of females after eclosion, intact and wingless by amputation. In A-D each datapoint represents data from a 10 min recording of one male with the indicated female, error bars indicate median and interquartile range.

## Discussion

### Immature female wing flicks act as signals of asexuality

We document wing flicking in immature females as a behavior that produces characteristic sound pulses that are distinct from other acoustic communication signals of *Drosophila melanogaster*, such as male courtship song or female copulation song (**Figure 1**). Wing flicks only occur when immatures are courted by mature males during short time window after eclosion (**Figure 2A**). In accordance with recent studies (Chen et al. 2022, Seong et al. 2023), we find that this time window up to approximately 10 h after eclosion represents a period of asexuality (**Figure 2B**). The probability of wing flicking sharply decreases 12 h after eclosion, when females start to accept matings and display an increasing receptivity. Wing flicks thus announce to the courting male the near certainty that there is no opportunity for copulation. They can be considered as an honest signal in mate choice, i.e. a signal that reliably displays the condition of the sender (Smith and Harper 2003). The response to an honest signal is predicted to not only benefit the sender, but also the receiver.

When exposed to wing flicking immatures, courting males produce shorter pulse song bouts and almost never attempt copulation (**Figure 4B, C**). Suppression of high-level mating effort directed at unreceptive targets might save energy and allow the male to switch to other, more promising targets when flies are in larger group. Attempted copulation appears to incur a particularly high cost, since the male shuts off processing of threats when he transitions to such a late state of courtship behavior (Cazalé-Debat et al. 2025), exposing himself and his object to the risk of predation.

For immature females, copulation attempts probably represent not only increased predation risk, but also a physical disturbance. Immature flies have not yet reached full ability to escape by flight (**Figure 2C**) and have a cuticle still undergoing final tanning and hardening (Ewers and White 2014), which might make them especially vulnerable to male harassment.

Immature wing flicks serve as a signal for asexuality but are not required to prevent copulations in the first 10-12 h time window after eclosion (**Figure 4E**). The proximate causes and adaptive value of immature asexuality are not completely understood. After eclosion, females have small ovaries that lack mature egg chambers (Manning 1967). Yolk accumulation (vitellogenesis) and maturation of fertilizable oocytes only start at the end of asexuality at around 12 h (Zhang et al. 2022). When lacking mature eggs, it might be disadvantageous for the female to invest in sperm storage. Some authors report rare forced matings between mature males and immature females with unexpanded wings within the first hour of eclosion in natural populations and under certain laboratory conditions (Markow 2000, Seeley and Dukas 2011), a phenomenon that we and others (Seong et al. 2023) could not observe. Interestingly, such teneral matings were found to be fertile, but produce around 70% less progeny than matings between mature individuals (Seeley and Dukas 2011), which could be an indication that sperm storage is not yet optimally developed in immature females.

In summary, immature wing flicking is a transient, developmentally tightly controlled signal that communicates asexuality and wards off males in a time period when mating is likely to be not yet of adaptive value to the female.

### State dependent rejection response to male courtship in immature versus mature mated females

While immature females respond to male courtship with wing flicking, mated females frequently extend their ovipositor (Connolly and Cook 1971, Kimura et al. 2015). How do wing flicking and ovipositor extrusion differ and why might females deploy two different behaviors correlated with lack of receptivity depending on their state?

As immature flicks (**Figure 3D**), ovipositor extrusion is triggered by male courtship and depends on courtship song. Full extrusion of the ovipositor prevents intromission and successful copulation, but loss of the behavior does not restore receptivity (Mezzera et al. 2020, Wang et al. 2020). In contrast to wing flicks (**Figure 4C**), ovipositor extrusion of mated females does not suppress copulation attempts but instead triggers them with high efficiency (Mezzera et al. 2020, Wang et al. 2020). This counterintuitive male response might be explained by the fact that mated females normally remate several times, a behavior that is clearly revealed when they are observed in a more natural setting in the presence of food and conspecifics over longer time periods (Lefevre and Jonsson 1962, Harshman and Clark 1998, Gorter et al. 2016).

Rather than completely shutting down receptivity, mated females become more selective in choosing a mate (Kohlmeier et al. 2021). Males achieving copulation with already mated females have an advantage for fertilization (Manier et al. 2010, Laturney et al. 2018) and might profit from the already stimulated fecundity of the female (Sirot et al. 2011).

For males, it therefore seems adaptive to probe a mated female after ovipositor extrusion with a copulation attempt, instead of depressing courtship. Attempts toward mated females might not only bring a chance for copulation but could also represent a form of genital sampling providing more detailed sensory information about the mating state and the presence of an egg or mating plug (von Philipsborn 2020).

In conclusion, the different rejection behaviors of immature and mated females elicit different responses in courting males, matching asexuality and selectively low receptivity of the females, respectively.

### The role of pC1a neurons in flicking behavior and receptivity

We find that inhibition of all Dsx+ or only pC1a neurons in immature virgins suppresses immature wing flicks (**Figure 3A**). This finding appears surprising, given the fact that inhibition of pC1a neurons in mature virgins suppresses receptivity (Yun et al. 2024). The observation that pC1a is required for rejection behavior in the immature state and for acceptance of copulation in the mature state suggests that the neuron is activated by courtship stimuli in both states, but that maturation changes processing in the circuits downstream of pC1a. This hypothesis contrasts with a model where the increase of sexual receptivity during maturation is linked to the cAMP dependent increase of pC1 excitability mediated by Dh44 (Kim et al. 2024).

In a larger group of 14 pC1 cells comprising different subtypes, isolated male courtship stimuli (pulse song or the application of male pheromone cVA) elicit less calcium transients in 1 d old virgins than in fully receptive 3 d old virgins (Kim et al. 2024). These data do not exclude the possibility that pC1a as a single pC1 subtype responds sufficiently to a combination of courtship stimuli in immature females to mediate immature flicking. Since flicking peaks only in a short time window of asexuality around 5 h after eclosion, transient pC1a excitability via an cAMP independent mechanism during this earlier phase could likewise explain our results.

Female receptivity is not yet at its maximum immediately after the “switch-on” event, which coincides with the rapid disappearance of flicking behavior between 10 - 15 h after eclosion but reaches a peak only at 3 d. In this longer phase of behavioral maturation between 18 h and 3 d, the comparatively low virgin female receptivity has been shown to also be due to the neuropeptide Leukokinine (LK), which inhibits pC1 neurons (Chen et al. 2022). While loss of LK receptor in pC1 neurons increases virgin receptivity at 36 h, it has no effect on receptivity in more immature virgins at 18 h (Chen et al. 2022), indicating the existence of alternative or additional mechanisms to prevent mating during the early phase of asexuality.

In summary, sexual maturation might progress through two phases: an early phase of asexuality, characterized by pC1a mediated wing flicking behavior of immature females in response to male courtship, and a subsequent prolonged “sexual transition”, during which receptivity increases while pC1 increases in excitability and activity due to a rise in DH44 and a decline in LK signals. In the future, uncovering the neuronal and molecular mechanism by which wing flicking behavior is lost will probe this hypothesis and give insight into the multilayered processes during sexual maturation.

### Relationship of immature female wing flicks and mature male aggression flicks

Mature males flick their wings during aggressive interactions when competing for resources such as a female or food (Jonsson et al. 2011, Versteven et al. 2017, Swain and von Philipsborn, Hindmarsh Sten et al. 2025). These male agonistic flicks produce pulse trains that resemble the ones produced by immature female flicks (**Figure 1A, D**). Aggression behavior is sexually dimorphic and in mature flies, wing flicking has only been reported in males (Nilsen et al. 2004). Like immature female wing flicks, male agonistic flicks toward a rival occur in response to courtship song and depend on integration of auditory and olfactory cues (Hinmarsh Sten et al. 2025). Immature female wing flicks suppresses copulation attempts toward immature females (**Figure 4C**). Male agonistic flicks have been hypothesized to physically hinder other males to mate with a female, as well as to acoustically interfere with the female hearing male courtship song, thereby momentarily decreasing copulation success in a competitive setting. When a male is exposed to wing flicks from a rival while he courts a female, his probability to be closely behind the female, i.e. in a position required for copulation attempts, decreases (Hindmarsh Sten et al. 2025).

The similar motor pattern, releasing stimuli and behavioral effects of immature female and male agonistic flicks raise the question if the two behavioral responses are controlled by the same neuronal circuits. pC1 neurons, which are required for immature female wing flicking (**Figure 3E**), are also required for male agonistic flicks. pC1 cells are a morphologically and functionally heterogenous group of neurons that are sex-specific or sexually dimorphic and depend in their total numbers and branching pattern on the sex determination factors Fruitless and Doublesex (Asahina 2018, Chen et al. 2025). We find that among the 5 described subtypes of the female pC1 cluster, only pC1a has an effect on immature female flicking (**Figure 3E**). The male equivalent of female pC1a, pC1x_a, differs in its branching pattern in the dorsal brain (Berg et al. 2025). For males, it has not yet been determined which subtype of pC1 neurons is involved in agonistic flicking, but pC1x_a is among the neurons manipulated in the study by Hindmarsh Sten et al. Male flicking also depends on a male specific neuron arborizing the wing neuropil of the ventral nerve cord (Hindmarsh Sten et al. 2025), which bears similarity to male specific neuronal class vPR6 that is required for male courtship song (von Philipsborn et al. 2011). This suggests that immature female and mature males effectuate flicking by sexually dimorphic circuits.

In some mating systems, one sex can mimic morphologically or behaviorally the other as a part of its reproductive strategy (Thornhill and Alcock 1983). For most demonstrated cases, e.g. in bluegill fish (Dominey 1981), salamander (Arnold 1976), rove beetle (Forsyth and Alcock 1990) or harvestman spider (Solano-Brenes et al. 2026), males that are at disadvantage in direct competition adopt a female mimicry to gain access to females guarded by a rival or disrupt a stronger male’s mating success. Do competing males in Drosophila mimic rejection signals of immature females or vice versa? We lack information how and in which order immature female and male agonistic wing flicking evolved. Possibly, the two behaviors developed in parallel, with the immature female rejection signal benefitting from male behavioral adaptation to male-male competition and the male agonistic signal becoming more effective through male behavioral adaptation to immature asexuality.

## Methods

### Drosophila culture

Drosophila melanogaster was cultured on medium made from cornmeal, oatmeal, sucrose and yeast in humidity (60-70%) and temperature (25°C) controlled incubators at a 12h:12h light dark cycle. As wild type (*wt*) strain, Canton S flies were used. All transgenic flies used in experiments are listed in Table 1.

**Table 1.** Transgenic flies used in experiment. Full genotypes corresponding to labels in the indicated figures and the stock origin/identifier of the transgenic stocks used for generating these genotypes. + denotes wild type chromosomes derived from wild type Canton S stock.

| Short name, label | Genotype | Figure | Transgenic stock origin, Identifier/RRID |
| --- | --- | --- | --- |
| *wt* | *+/+; +/+ Canton S wild type strain* | 1, 2, 3A-D, 4 | Gift from B. Dickson |
| *fruF* ctrl | *+/+; fruF/+* | 3D | RRID: BDSC_66873 |
| *fruF* | *+/+; fruF/fruFLP* | 3D | RRID: BDSC_66870 (*FruFLP*) |
| *anosmic* | *IR8a<sup>1</sup>/ IR8a; IR25a<sup>2</sup>/IR25a<sup>2</sup>; GR63a<sup>1</sup>, Orco<sup>1</sup>/ GR63a<sup>1</sup>, Orco<sup>1</sup>* | 3D | Gift from R. Benton, Ramdya et al. 2015 |
| *fruM* | *+/+; fruM/+* | 3D | RRID: BDSC_66874 |
| *Dsx* ctrl | *+/+; Dsx-GAL4/+* | 3E, 4A-C | RRID: BDSC_66674 |
| *TNT* ctrl | *+/+; UAS-TNT, UAS-mCD8-Egfp/+; +/+* | 3E, 4A-C | Gift from B. Dickson |
| *Dsx > TNT* | *+/+; UAS-TNT, UAS-mCD8-Egfp/+; Dsx-GAL4/+* | 3E, 4A-C |  |
| *pC1a* split ctrl | *+/+; R52G04-p65.AD.attp40/+;* | 3E | RRID: BDSC_71085 |
| *TNT Dsx* split ctrl | *+/+; UAS-TNT, UAS-mCD8-Egfp/+; Dsx-DBD/+* | 3E | RRID: BDSC_89697 (*Dsx-DBD*) |
| *pC1a > TNT* | *+/+; UAS-TNT, UAS-mCD8-Egfp/ R52G04-p65.AD.attp40; Dsx-DBD/+* | 3E |  |
| *pC1a > Egfp* | *+/+; UAS-mCD8-Egfp/ R52G04-p65.AD.attp40; Dsx-DBD/+* | 3E |  |

### Behavioral assays

Immature females were collected on light CO_2_ anesthesia after eclosion as soon as wing expansion was complete, male flies were collected within 4 h of eclosion. After collection, flies were aged in isolation in flat bottom 1.5 ml 96 well blocks filled with 0.5 ml food per well and covered with a PCR foil with air holes (von Philipsborn et al. 2023a). For obtaining reproductively mature males or females, flies were aged for 4-7 days. For obtaining mature mated females, 4 d old virgin females were individually paired with a male in small chambers and recollected after the end of copulation.

Fly wing sounds were recorded at a sampling rate of 10 kHz with a custom-built multi-channel microphone array in chambers of 1 cm diameter, amplified and digitized as described previously (von Philipsborn et al. 2023b, Arthur et al. 2013). Immature female wing flicks were recorded by pairing a mature male with an immature female. Mature male agonistic wing flicks were recorded by pairing two mature males in the presence of a mature virgin. High speed recordings of wing movements were acquired by an Optronis CR3000x2, monochrome camera with a Sigma 105mm f2.8 macro lens. For assessing the cumulative mating rate after eclosion, female pupae were placed with a single mature male in a 35 ml food vial with a sloped surface and imaged after eclosion by time lapse every 2 s over 40 h with consumer camcorders (JVC Everio) (von Philipsborn et al. 2023c). Forced flight tests were performed as previously (Loreau et al. 2025), by dropping flies in a 1 m high plexiglass cylinder with 5 marked landing zones along its height. For testing the frequency of copulation attempts, males and females were videotaped with consumer camcorders in standard courtship chambers of 1 cm diameter.

### Precocene I treatment

Precocene treatment was performed as described before (Ringo et al. 2005). 2.2 μmol precocene I (7-methoxy-2,2-dimethyl chromene, Santa Cruz Biotechnologies, CAS: 17598-02-6) was dissolved in acetone, applied on a 1 x 2 cm piece of filter paper and air dried. Cumulative mating after eclosion was tested in presence of the precocene I paper in the food vial. Wing flicking after precocene I treatment was tested after housing 10 females for 24 h in the presence of a precocene I paper in a 23 ml food vial. As a control, acetone treated paper was used.

### Amputation experiments

For wing and arista amputation experiments, body parts were cut under light CO_2_ anesthesia with Vanna’s spring scissors (Fine science tools). As control, mock amputated animals handled under CO_2_ anesthesia were used.

### Immunohistochemistry and confocal imaging

Immunohistochemistry of the central nervous system and confocal was performed as described previously before (O’Sullivan et al. 2018). After dissection and fixation in 4% paraformaldehyde, tissues were stained with mouse nc82 antibody (Developmental Studies Hybridoma Bank, RRID:AB_2314866), rabbit anti-GFP antibody (Torrey Pines Biolabs, RRID: AB_10013661) followed by respective secondary antibodies (Goat IgG (H+L) Alexa Fluor 488 and 647) and mounted in Vectashield (Vector labs, RRID: AB_2336789). Confocal image stacks were acquired at a Leica STELLARIS8 FALCON microscope equipped with a HC PL APO 20x/0,75 multi-immersion objective.

### Quantification and data analysis

For statistical tests, Graph Pad prism software and R software were used. Tests and outcomes, sample size and further details of what each datapoint represents can be found in the figure legends. Acoustic recordings were analyzed with a previously reported MATLAB script (von Philipsborn et al. 2023b, Arthur et al. 2013) and by visualization of oscillograms to manually correct automated detection of pulse events. For assessment of IPI and train length, pulses spaced at 15 ms-150 ms for male courtship song and at 15 ms-300 ms for immature female or matrue male wing flicks were considered to be part of one train. For analyzing the latencies between male song and immature flicks, we measured values for each immature flick trains from 42 pairs recorded for 10 min, pooled the data and binned it in 100 ms wide bins. To perform a permutation test, acoustic signals from each female were virtually aligned with courtship song of all 41 males that were not the real partner of the female, resulting in 1722 false pairings. For a permuted dataset we randomly selected 42 unique false pairs from this pool. In Figure 3C, mean probabilities of 1000 permuted datasets are plotted. Permutation test p values were calculated generating 10.000 permuted datasets, asking the frequency of observing a flick occurrence probability of at least as high as seen in the real dataset in a given latency bin.

## Acknowledgements

We thank Valerie Salicio, Laurence Clément, Isabelle Scerri and David Rodriguez Crespo for technical assistance and Rachel Ververidis for help with data quantification and analysis. We thank the DGRC stock library, Richard Benton and Barry Dickson for providing fly stocks, Developmental Studies Hybridoma Bank for the nc82 antibody, and the University of Fribourg Bioimaging Core Facility for assistance with microscopy. This study was supported by Swiss National Science foundation grant 310030_212222.

## Notes

### Competing Interest Statement

The authors have declared no competing interest.

## References

Allen, A.M., Neville, M.C., Nojima, T., Alejevski, F., and Goodwin, S.F. (2026). Differential neuronal survival defines a novel axis of sexual dimorphism in the Drosophila brain. Cell Genomics 6, 101125.

Arnold, A.P. (2009). The organizational–activational hypothesis as the foundation for a unified theory of sexual differentiation of all mammalian tissues. Hormones and Behavior 55, 570–578.

Arnold, S.J. (1976). Sexual Behavior, Sexual Interference and Sexual Defense in the Salamanders Ambystoma maculatum, Ambystoma tigrinum and Plethodon jordani. Zeitschrift für Tierpsychologie 42, 247–300.

Arthur, B., Sunayama-Morita, T., Coen, P., Murthy, M., and Stern, D. (2013). Multi-channel acoustic recording and automated analysis of Drosophila courtship songs. BMC Biology 11, 11.

Asahina, K. (2018). Sex differences in Drosophila behavior: qualitative and quantitative dimorphism. Current Opinion in Physiology 6, 35–45.

Baker, C.A., Clemens, J., and Murthy, M. (2019). Acoustic Pattern Recognition and Courtship Songs: Insights from Insects. Annual Review of Neuroscience 42, 129–147.

Berg, S., Beckett, I.R., Costa, M., Schlegel, P., Januszewski, M., Marin, E.C., Nern, A., Preibisch, S., Qiu, W., Takemura, S.-y., et al. (2025). Sexual dimorphism in the complete connectome of the Drosophila male central nervous system. bioRxiv, 2025.2010.2009.680999.

Bilen, J., Atallah, J., Azanchi, R., Levine, J.D., and Riddiford, L.M. (2013). Regulation of onset of female mating and sex pheromone production by juvenile hormone in Drosophila melanogaster. Proceedings of the National Academy of Sciences 110, 18321–18326.

Bownes, M. (1982). Hormonal and Genetic Regulation of Vitellogenesis in Drosophila. The Quarterly Review of Biology 57, 247–274.

Bownes, M., and Rembold, H. (1987). The titre of juvenile hormone during the pupal and adult stages of the life cycle of Drosophila melanogaster. European Journal of Biochemistry 164, 709–712.

Box, A.M., Ramesh, N.A., Nandakumar, S., Church, S.J., Prasad, D., Afrakhteh, A., Taichman, R.S., and Buttitta, L. (2024). Cell cycle variants during Drosophila male accessory gland development. G3 Genes|Genomes|Genetics 14.

Cazalé-Debat, L., Scheunemann, L., Day, M., Fernandez-d.V. Alquicira, T., Dimtsi, A., Zhang, Y., Blackburn, L.A., Ballardini, C., Greenin-Whitehead, K., Reynolds, E., et al. (2024). Mating proximity blinds threat perception. Nature 634, 635–643.

Chen, J., Tu, W., Li, Z., Ma, M., Jiang, S., Guan, W., Wang, R., Pan, Y., and Peng, Q. (2025). Diverse functions of sex determination gene doublesex on sexually dimorphic neuronal development and behaviors. Journal of Genetics and Genomics 52, 1199–1210.

Chen, J., Zhu, P., Jin, S., Zhang, Z., Jiang, S., Li, S., Liu, S., Peng, Q., and Pan, Y. (2025). A hormone-to-neuropeptide pathway inhibits sexual receptivity in immature Drosophila females. Proceedings of the National Academy of Sciences 122, e2418481122.

Connolly, K., and Cook, R. (1973). Rejection Responses by Female Drosophila melanogaster: Their Ontogeny, Causality and Effects upon the Behaviour of the Courting Male. Behaviour 44, 142–166.

Day, M., and Rezaval, C. (2026). Neural basis of sexually dimorphic decision-making. Current Opinion in Neurobiology 96, 103146.

Demir, E., and Dickson, B.J. (2005). fruitless Splicing Specifies Male Courtship Behavior in Drosophila. Cell 121, 785–794.

Dominey, W.J. (1981). Maintenance of female mimicry as a reproductive strategy in bluegill sunfish (Lepomis macrochirus). Environmental Biology of Fishes 6, 59–64.

Ellendersen, B.E., and von Philipsborn, A.C. (2017). Neuronal modulation of D. melanogaster sexual behaviour. Current Opinion in Insect Science 24, 21–28.

Ewing, A.W., and Bennet-Clark, H.C. (1968). The Courtship Songs of Drosophila. Behaviour 31, 288–301.

Feng, K., Palfreyman, Mark T., Häsemeyer, M., Talsma, A., and Dickson, Barry J. (2014). Ascending SAG Neurons Control Sexual Receptivity of Drosophila Females. Neuron 83, 135–148.

Ford, S.C., Napolitano, L.M., McRobert, S.P., and Tompkins, L. (1989). Development of behavioral competence in young Drosophila melanogaster adults. Journal of Insect Behavior 2, 575–588.

Forsyth, A., and Alcock, J. (1990). Female mimicry and resource defense polygyny by males of a tropical rove beetle, Leistotrophus versicolor (Coleoptera : Staphylinidae). Behavioral Ecology and Sociobiology 26, 325–330.

Goodwin, S.F., and Hobert, O. (2021). Molecular Mechanisms of Sexually Dimorphic Nervous System Patterning in Flies and Worms. Annual Review of Cell and Developmental Biology 37, 519–547.

Gorter, J.A., Jagadeesh, S., Gahr, C., Boonekamp, J.J., Levine, J.D., and Billeter, J.-C. (2016). The nutritional and hedonic value of food modulate sexual receptivity in Drosophila melanogaster females. Scientific Reports 6, 19441.

Hadjaje, S., Andrade-Silva, I., Dalbe, M.-J., Clément, R., and Marthelot, J. (2024). Mechanics of Drosophila wing deployment. Nature Communications 15, 10577.

Harshman, L.G., and Clark, A.G. (1998). Inference of sperm competition from broods of dield-caught Drosophila. Evolution 52, 1334–1341.

Häsemeyer, M., Yapici, N., Heberlein, U., and Dickson, B.J. (2009). Sensory Neurons in the Drosophila Genital Tract Regulate Female Reproductive Behavior. Neuron 61, 511–518.

Herbison, A.E. (2016). Control of puberty onset and fertility by gonadotropin-releasing hormone neurons. Nature Reviews Endocrinology 12, 452–466.

Herndon, L.A., Chapman, T., Kalb, J.M., Lewin, S., Partridge, L., and Wolfner, M.F. (1997). Mating and hormonal triggers regulate accessory gland gene expression in male Drosophila. Journal of Insect Physiology 43, 1117–1123.

Hindmarsh Sten, T., Li, R., Hollunder, F., Eleazer, S., and Ruta, V. (2025). Male-male interactions shape mate selection in Drosophila. Cell 188, 1486–1503.e1425.

Ji, X., Li, X., Wang, L., Liu, S., Jiang, X., and Pan, Y. (2023). Asexuality in Drosophila juvenile males is organizational and independent of juvenile hormone. The EMBO Reports 24, EMBR202356898.

Jonsson, T., Kravitz, E.A., and Heinrich, R. (2011). Sound production during agonistic behavior of male Drosophila melanogaster. Fly 5, 29–38.

Karigo, T., and Deutsch, D. (2022). Flexibility of neural circuits regulating mating behaviors in mice and flies. Frontiers in Neural Circuits Volume 16 -2022.

Kerwin, P., Yuan, J., and von Philipsborn, A.C. (2020). Female copulation song is modulated by seminal fluid. Nature Communications 11, 1430.

Kim, D.-H., Jang, Y.-H., Yun, M., Lee, K.-M., and Kim, Y.-J. (2024). Long-term neuropeptide modulation of female sexual drive via the TRP channel in Drosophila melanogaster. Proceedings of the National Academy of Sciences 121, e2310841121.

Kimura, K.-I. (2011). Role of cell death in the formation of sexual dimorphism in the Drosophila central nervous system. Development, Growth & Differentiation 53, 236–244.

Kimura, K.-i., Sato, C., Koganezawa, M., and Yamamoto, D. (2015). Drosophila Ovipositor Extension in Mating Behavior and Egg Deposition Involves Distinct Sets of Brain Interneurons. PLOS ONE 10, e0126445.

Kohlmeier, P., Zhang, Y., Gorter, J.A., Su, C.-Y., and Billeter, J.-C. (2021). Mating increases Drosophila melanogaster females’ choosiness by reducing olfactory sensitivity to a male pheromone. Nature Ecology & Evolution.

Laturney, M., van Eijk, R., and Billeter, J.-C. (2018). Last male sperm precedence is modulated by female remating rate in Drosophila melanogaster. Evolution Letters 2, 180–189.

Lefevre, G., and Jonsson, U.B. (1962). Sperm transfer, storage, displacement, and utilization in Drosophila melanogaster. Genetics 47, 1719–1736.

Lin, H.-H., Cao, D.-S., Sethi, S., Zeng, Z., Chin, Jacqueline S.R., Chakraborty, Tuhin S., Shepherd, Andrew K., Nguyen, Christine A., Yew, Joanne Y., Su, C.-Y., et al. (2016). Hormonal Modulation of Pheromone Detection Enhances Male Courtship Success. Neuron 90, 1272–1285.

Liu, H., and Kubli, E. (2003). From the Cover: Sex-peptide is the molecular basis of the sperm effect in Drosophila melanogaster. Proceedings of the National Academy of Sciences 100, 9929–9933.

Loreau, V., Koolhaas, W.H., Chan, E.H., De Boissier, P., Brouilly, N., Avosani, S., Sane, A., Pitaval, C., Reiter, S., Luis, N.M., et al. (2025). Titin-dependent biomechanical feedback tailors sarcomeres to specialized muscle functions in insects. Science Advances 11, eads8716.

Manier, M.K., Belote, J.M., Berben, K.S., Novikov, D., Stuart, W.T., and Pitnick, S. (2010). Resolving Mechanisms of Competitive Fertilization Success in Drosophila melanogaster. Science 328, 354–357.

Manning, A. (1967). The control of sexual receptivity in female Drosophila. Anim Behav 15, 239–250.

Markow, T.A. (2000). Forced Matings in Natural Populations of Drosophila. The American Naturalist 156, 100–103.

McCarthy, M.M., and Arnold, A.P. (2011). Reframing sexual differentiation of the brain. Nature Neuroscience 14, 677–683.

Meeh, K.L., Rickel, C.T., Sansano, A.J., and Shirangi, T.R. (2021). The development of sex differences in the nervous system and behavior of flies, worms, and rodents. Developmental Biology 472, 75–84.

Meiselman, M., Lee, S.S., Tran, R.-T., Dai, H., Ding, Y., Rivera-Perez, C., Wijesekera, T.P., Dauwalder, B., Noriega, F.G., and Adams, M.E. (2017). Endocrine network essential for reproductive success in Drosophila melanogaster.Proceedings of the National Academy of Sciences 114, E3849–E3858.

Mezzera, C., Brotas, M., Gaspar, M., Pavlou, H.J., Goodwin, S.F., and Vasconcelos, M.L. (2020). Ovipositor Extrusion Promotes the Transition from Courtship to Copulation and Signals Female Acceptance in Drosophila melanogaster. Current Biology 30, 3736–3748.e3735.

Moshitzky, P., Fleischmann, I., Chaimov, N., Saudan, P., Klauser, S., Kubli, E., and Applebaum, S.W. (1996). Sex-peptide activates juvenile hormone biosynthesis in the Drosophila melanogaster corpus allatum. Archives of Insect Biochemistry and Physiology 32, 363–374.

Nilsen, S.P., Chan, Y.-B., Huber, R., and Kravitz, E.A. (2004). Gender-selective patterns of aggressive behavior in Drosophila melanogaster. Proceedings of the National Academy of Sciences of the United States of America 101, 12342–12347.

O’Sullivan, A., Lindsay, T., Prudnikova, A., Erdi, B., Dickinson, M., and von Philipsborn, A.C. (2018). Multifunctional Wing Motor Control of Song and Flight. Current Biology 28, 13.

Pailette, M., Ikeda, H., and Jallon, J.M. (1991). A new acoustic Signal of the fruit-flies Drosophila simulans and D. melanogaster. Bioacoustics 3, 247–254.

Peng, J., Chen, S., Büsser, S., Liu, H., Honegger, T., and Kubli, E. (2005). Gradual Release of Sperm Bound Sex-Peptide Controls Female Postmating Behavior in Drosophila. Current Biology 15, 207–213.

Ramdya, P., Lichocki, P., Cruchet, S., Frisch, L., Tse, W., Floreano, D., and Benton, R. (2014). Mechanosensory interactions drive collective behaviour in Drosophila. Nature 519, 233.

Ramesh, N.A., Box, A.M., and Buttitta, L.A. (2025). Post-eclosion growth in the Drosophila ejaculatory duct is driven by Juvenile hormone signaling and is essential for male fertility. Developmental Biology 519, 122–141.

Ringo, J., Talyn, B., and Brannan, M. (2005). Effects of Precocene and Low Protein Diet on Reproductive Behavior in Drosophila melanogaster (Diptera: Drosophilidae). Annals of the Entomological Society of America 98, 601–607.

Ringo, J., Werczberger, R., Altaratz, M., and Segal, D. (1991). Female sexual receptivity is defective in juvenile hormone-deficient mutants of theapterous gene of Drosophila melanogaster. Behavior Genetics 21, 453–469.

Santos, C.G., Humann, F.C., and Hartfelder, K. (2019). Juvenile hormone signaling in insect oogenesis. Current Opinion in Insect Science 31, 43–48.

Sato, K., and Yamamoto, D. (2020). The mode of action of Fruitless: Is it an easy matter to switch the sex? Genes, Brain and Behavior 19, e12606.

Schretter, C.E., Aso, Y., Robie, A.A., Dreher, M., Dolan, M.J., Chen, N., Ito, M., Yang, T., Parekh, R., Branson, K.M., et al. (2020). Cell types and neuronal circuitry underlying female aggression in Drosophila. Elife 9.

Schulz, K.M., and Sisk, C.L. (2016). The organizing actions of adolescent gonadal steroid hormones on brain and behavioral development. Neuroscience & Biobehavioral Reviews 70, 148–158.

Seeley, C., and Dukas, R. (2011). Teneral matings in fruit flies: male coercion and female response. Animal Behaviour 81, 595–601.

Seong, K.-H., Uemura, T., and Kang, S. (2023). Road to sexual maturity: Behavioral event schedule from eclosion to first mating in each sex of Drosophila melanogaster. iScience 26, 107502.

Shao, L., Chung, P., Wong, A., Siwanowicz, I., Kent, C.F., Long, X., and Heberlein, U. (2019). A Neural Circuit Encoding the Experience of Copulation in Female Drosophila. Neuron 102, 1025–1036.e1026.

Singh, A., Buehner, N.A., Lin, H., Baranowski, K.J., Findlay, G.D., and Wolfner, M.F. (2018). Long-term interaction between Drosophila sperm and sex peptide is mediated by other seminal proteins that bind only transiently to sperm. Insect Biochemistry and Molecular Biology 102, 43–51.

Sirot, L.K., Wolfner, M.F., and Wigby, S. (2011). Protein-specific manipulation of ejaculate composition in response to female mating status in Drosophila melanogaster. Proceedings of the National Academy of Sciences 108, 9922–9926.

Smith, J.M., and Harper, D. (2003). Animal Signals, (Oxford University Press).

Solano-Brenes, D., Furlan, C.M., da Silva, R.C., Willemart, R.H., Nascimento, F.S., and Machado, G. (2026). Same-sex sexual behavior in a male-dimorphic arachnid with a resource-defense polygyny: evidence for female mimicry? The Science of Nature 113, 8.

Soller, M., Bownes, M., and Kubli, E. (1999). Control of Oocyte Maturation in Sexually Mature Drosophila Females. Developmental Biology 208, 337–351.

Sugime, Y., Watanabe, D., Yasuno, Y., Shinada, T., Miura, T., and Tanaka, N.K. (2017). Upregulation of Juvenile Hormone Titers in Female Drosophila melanogaster Through Mating Experiences and Host Food Occupied by Eggs and Larvae. Zoological Science 34, 52–57, 56.

Swain, B., and von Philipsborn, A.C. (2021). Chapter Three -Sound production in Drosophila melanogaster: Behaviour and neurobiology. In Advances in Insect Physiology, Volume 61, R. Jurenka, ed. (Academic Press), pp. 141–187.

Thornhill, R., and Alcock, J. (1983). The evolution of insect mating systems, (Cambridge, Mass: Harvard University Press).

Tong, X.-J., Wang, F., and Xu, X. (2026). Sexually dimorphic neural circuits underlying mating behaviors: Insights from worms, flies, and mice. Current Opinion in Neurobiology 96, 103151.

Truman, J.W., and Riddiford, L.M. (2023). Drosophila postembryonic nervous system development: a model for the endocrine control of development. Genetics 223.

Versteven, M., Vanden Broeck, L., Geurten, B., Zwarts, L., Decraecker, L., Beelen, M., Göpfert, M.C., Heinrich, R., and Callaerts, P. (2017). Hearing regulates Drosophila aggression. Proceedings of the National Academy of Sciences 114, 1958–1963.

von Philipsborn, A.C. (2020). Neuroscience: The Female Art of Saying No. Current Biology 30, R1080–R1083.

von Philipsborn, A.C., Liu, T., Yu, J.Y., Masser, C., Bidaye, S.S., and Dickson, B.J. (2011). Neuronal Control of Drosophila Courtship Song. Neuron 69, 509–522.

von Philipsborn, A.C., Shohat-Ophir, G., and Rezaval, C. (2023a). Single-Pair Courtship and Competition Assays in Drosophila. Cold Spring Harbor Protocols 2023, pdb.prot108105.

von Philipsborn, A.C., Shohat-Ophir, G., and Rezaval, C. (2023b). Probing Acoustic Communication during Fly Reproductive Behaviors. Cold Spring Harbor Protocols 2023, pdb.prot108107.

von Philipsborn, A.C., Shohat-Ophir, G., and Rezaval, C. (2023c). Female Fly Postmating Behaviors. Cold Spring Harbor Protocols 2023, pdb.prot108108.

Wang, F., Wang, K., Forknall, N., Parekh, R., and Dickson, B.J. (2020a). Circuit and Behavioral Mechanisms of Sexual Rejection by Drosophila Females. Current Biology 30, 3749–3760.e3743.

Wang, F., Wang, K., Forknall, N., Patrick, C., Yang, T., Parekh, R., Bock, D., and Dickson, B.J. (2020b). Neural circuitry linking mating and egg laying in Drosophila females. Nature 579, 101–105.

Wang, K., Wang, F., Forknall, N., Yang, T., Patrick, C., Parekh, R., and Dickson, B.J. (2021). Neural circuit mechanisms of sexual receptivity in Drosophila females. Nature 589, 577–581.

White, B.H., and Ewer, J. (2014). Neural and Hormonal Control of Postecdysial Behaviors in Insects. Annual Review of Entomology 59, 363–381.

Wijesekera, T.P., Saurabh, S., and Dauwalder, B. (2016). Juvenile Hormone Is Required in Adult Males for Drosophila Courtship. PLOS ONE 11, e0151912.

Wilson, T.G., DeMoor, S., and Lei, J. (2003). Juvenile hormone involvement in Drosophila melanogaster male reproduction as suggested by the Methoprene-tolerant27 mutant phenotype. Insect Biochemistry and Molecular Biology 33, 1167–1175.

Yang, C.-h., Rumpf, S., Xiang, Y., Gordon, M.D., Song, W., Jan, L.Y., and Jan, Y.-N. (2009). Control of the Postmating Behavioral Switch in Drosophila Females by Internal Sensory Neurons. Neuron 61, 519–526.

Yapici, N., Kim, Y.J., Ribeiro, C., and Dickson, B.J. (2008). A receptor that mediates the post-mating switch in Drosophila reproductive behaviour. Nature 451, 33–37.

Yip, C., Wyler, S.C., Liang, K., Yamazaki, S., Cobb, T., Safdar, M., Metai, A., Merchant, W., Wessells, R., Rothenfluh, A., et al. (2024). Neuronal E93 is required for adaptation to adult metabolism and behavior. Molecular Metabolism 84, 101939.

Yun, M., Kim, D.-H., Ha, T.S., Lee, K.-M., Park, E., Knaden, M., Hansson, B.S., and Kim, Y.-J. (2024). Male cuticular pheromones stimulate removal of the mating plug and promote re-mating through pC1 neurons in Drosophila females. eLife 13, RP96013.

Zhang, C., Kim, A.J., Rivera-Perez, C., Noriega, F.G., and Kim, Y.-J. (2022). The insect somatostatin pathway gates vitellogenesis progression during reproductive maturation and the post-mating response. Nature Communications 13, 969.

Zhang, S.X., Glantz, E.H., Miner, L.E., Rogulja, D., and Crickmore, M.A. (2021). Hormonal control of motivational circuitry orchestrates the transition to sexuality in Drosophila. Science Advances 7, eabg6926.

Zhou, C., Pan, Y., Robinett, Carmen C., Meissner, Geoffrey W., and Baker, Bruce S. (2014). Central Brain Neurons Expressing doublesex Regulate Female Receptivity in Drosophila. Neuron 83, 149–163.

